# Transfer learning framework via Bayesian group factor analysis incorporating feature-wise dependencies

**DOI:** 10.1101/2025.05.07.648613

**Authors:** Dharani Thirumalaisamy, Natasha Black, Mehmet Gönen, Olga Nikolova

## Abstract

Transfer learning considers distinct but related tasks defined over heterogeneous domains and aims to improve generalization and performance through knowledge transfer between tasks. This approach can be especially advantageous in biomedical contexts with insufficient labeled training data, where joint learning across domains can enable inference in otherwise underpowered datasets. High-dimensional biomedical data is characterized with redundancy, rendering non-linear dependencies among features. Existing models often fail to leverage such feature dependencies during inference, limiting their ability to model complex biological systems. We present a Bayesian group factor analysis transfer learning framework that supports multitask, multi-modal learning. Our approach learns a shared latent space within each domain, simultaneously across multiple domains, and uses a feature-wise prior to model complex relationships. We evaluate our framework using controlled synthetic data experiments and four disjoint patient cancer datasets from acute myeloid leukemia and neuroblastoma. We show that our method improves drug response prediction and more readily recapitulates consensus biomarkers of drug response. Similarly, our approach improves tumor purity prediction and identifies a robust gene signature associated with it. Our framework is scalable, interpretable, and adaptable across target phenotypes, offering a robust solution for a wide range of heterogeneous multi-omics problems.

## 1. Introduction

### 1.1 Previous work

Transfer learning has gained traction in cancer research, especially in leveraging knowledge from well-annotated datasets to improve performance on smaller or underpowered cohorts. However, most existing approaches are designed for specific settings or classification tasks (Karbalayghareh et al., 2018a; Hajiramezanali et al., 2018), with fewer models tailored to regression tasks.

Some supervised transfer learning models have addressed regression problems in cancer research, particularly in predicting drug responses. These approaches, including cost optimization and domain transfer learning techniques (Dhruba et al., 2018), often involve training on one cancer cohort (e.g., The Cancer Cell Line Encyclopedia (CCLE) (Barretina et al., 2012)) and predicting on another (e.g., The Genomics of Drug Sensitivity in Cancer (GDSC) (Yang et al., 2012)). However, these models typically rely on shared drug feature spaces and matched cell lines across cohorts, limiting their ability to generalize to disjoint patient cohorts.

More recently, deep learning based transfer learning approaches have been applied for drug response prediction (Baptista et al., 2021; Ju et al., 2025). These models are often trained on large pan-cancer datasets and fine-tuned on a specific cancer type or a drug. Although these models can achieve strong predictive performance, they often lack interpretability. While post-hoc interpretability methods such as SHAP (Lundberg and Lee, 2017) and LIME (Ribeiro et al., 2016) exist, they are not always reliable (Slack et al., 2020; Van den Broeck et al., 2022).

An ensemble-based deep learning transfer learning approach has also been explored to predict anti-cancer drug responses (Zhu et al., 2020). In this approach, models are trained on one cell line dataset and tested on another, using resources such as The Cancer Therapeutics Response Portal (CTRP) (Basu et al., 2013; Seashore-Ludlow et al., 2015; Rees et al., 2016), GDSC, CCLE and The Genentech Cell Line Screening Initiative (gCSI) (Haverty et al., 2016). However, this approach does not take dependencies between genomic features into account, which are known to interact through complex regulatory networks. In addition, it has not been applied to data beyond cell lines and code was not readily available for benchmarking in our study.

In parallel, fully Bayesian approaches for transfer learning have been proposed for various regression problems (Karbalayghareh et al., 2018b). These models leverage a joint prior distribution to transfer knowledge between the source and the target domain. However, they do not account for feature interactions or interpretability in the context of biological data.

Complementary to supervised strategies, several unsupervised methods have been developed to integrate multi-omics data, with Multi-Omics Factor Analysis (MOFA+) (Argelaguet et al., 2020) emerging as one of the most widely adopted approaches. These models effectively learn low-dimensional representations from heterogeneous data by capturing sources of variation across multiple modalities. However, these models do not account for domain shifts between source and target datasets. While MOFA+ can be used to identify predictive biomarkers, it cannot be used for prediction.

### 1.2 Contributions

We previously demonstrated that using a gene-wise prior to integrate genomic data improves drug response prediction and gene signature identification (Nikolova et al., 2017). The model employs a multitask Bayesian group factor analysis model (GBGFA) with a feature-wise prior that prioritizes genes that corroborate information in multiple measurement platforms. In subsequent work, we extended the use of this feature-wise prior in not only integrating molecular modalities, but also, for the modeling of drugs that have been screened in multiple datasets (Thirumalaisamy et al., 2024). We showed that a gene-wise prior, when combined with a drug-wise prior, recovers relevant biological signal even in the cases with conflicting drug response measurements, leading to more generalizable downstream associations. Building on this foundation, we now introduce a novel generative transfer learning framework that uses the feature-wise prior to simultaneously transfer information across cohorts. Like GBGFA, the model uses multitask multi-modal learning to find a shared latent space within and across cohorts, which enables the model to accommodate cohort-specific distribution shifts.

Here we derive a variational approximation inference algorithm that simultaneously solves multiple regression tasks by leveraging similarities in molecular and functional profiles across datasets. Additionally, the feature-wise priors enable us to trace which latent factors influence predictions, and which are the respective contributing features, making the model interpretable and accounting for feature-wise dependencies.

## 2. Materials

We evaluate our model’s performance on synthetic and primary cancer patient cohorts in acute myeloid leukemia (AML) and neuroblastoma (NB).

### 2.1 Synthetic data

Here we simulated data using our generative model to evaluate if the model effectively encodes our underlying assumptions. For complete notation please see section 3.1. We generated *C* = 2 cohorts, each with *M*_1_ = *M*_2_ = 2 views and *N*_*c*_ = 100 samples, where the first view had *D*_11_ = *D*_21_ = 200 and the second had *D*_12_ = *D*_22_ = 100 features, respectively. For each cohort, the latent space matrices 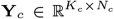 and the factor loadings 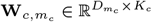 were sampled from a normal distribution using the `rnorm` R package, where 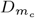 and *K*_*c*_ are the number of features and latent components, respectively. The observation matrices 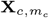 were generated for each view in every cohort by multiplying **Y**_*c*_ and 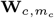, followed by the addition of Gaussian noise 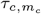 with mean 0 and variance 0.1 generated via the rnorm R package. When generating this data, the model is initialized with values of 1*e* − 14 for *α*_*γ*_, *β*_*γ*_, *α*_*τ*_ and *β*_*τ*_, with *α* and *β* being the shape and the scale hyperparameters of *γ* distribution, respectively. Training is performed with *K*_*c*_ = 50 latent components.

### 2.2 BeatAML acute myeloid leukemia patient cohort (BAML)

BeatAML is the largest assembled dataset of matched multi-omic profiles, drug screens, and clinical information for primary leukemia samples from the Beat AML project at the Knight Cancer Institute at OHSU (Tyner et al., 2018; Bottomly et al., 2022). Here we briefly summarize the data generating platforms used in this study. Gene expression data were acquired using the Illumina HiSeq 2500 platform and the measurements were reported as reads per kilobase per million mapped reads (RPKM). These RPKM values for protein coding genes were converted to transcripts per million (TPM) using Kallisto (Bray et al., 2016). The TPM gene expression data were *log*2 transformed with pseudo-count of 1. Mutation data were obtained using the Illumina Nextera RapidCapture Exome capture probes. The data was summarized by assigning a value of 1 to indicate the presence of a mutation and 0 otherwise. Drug response data for 164 drugs were collected via *ex vivo* functional drug screens on freshly isolated mononuclear patient cells. Drug response was quantified using the area under the drug dose–response curve (*AUC*), where higher values indicate resistance and lower – sensitivity to the respective compound. Low variance genes (*σ* < 0.001) and missing values were excluded from analysis.

### 2.3 Acute myeloid leukemia patient cohort from the Institute for Molecular Medicine Finland (FIMM)

Similarly to BeatAML, the FIMM dataset includes gene expression, mutation and *ex vivo* drug response primary patient data (Malani et al., 2022). Here we briefly summarize the data generating platforms used in this study. Gene expression and mutation data were acquired using Illumina HiSeq 1500, 2000, or 2500 instruments. Raw counts data for gene expression were transformed to TPM values using the convertCounts R package (Law et al., 2018). The TPM gene expression data were *log*2 transformed with pseudo-count of 1. *Ex vivo* drug response was measured for 538 emerging and FDA approved cancer drugs. Drug response in this dataset was reported as drug sensitivity score (*DSS*), which quantifies the relative inhibition of the cancer cells across varying drug concentrations. To ensure consistency with BAML’s *AUC* format, *DSS* values were transformed by subtracting each value from the maximum observed DSS, such that high values correspond to resistance, and low values indicate sensitivity, similar to the BeatAML *AUC*. Low variance genes (*σ* < 0.001) and missing values were similarly excluded.

### 2.4 Drug families and consensus biomarkers

We have curated drug information as it pertains to drugs screened in the BAML and FIMM patients cohorts. We use Supplementary File 4 from Bottomly et al (Bottomly et al., 2022) as a reference for drug families. Starting with the civicDB public database (Krysiak et al., 2023) and through literature examination, we have curated a list of consensus biomarkers of drug response (Supp. table ST1).

### 2.5 Neuroblastoma patient cohort from the Therapeutically Applicable Research to Generate Effective Treatments (TARGET) study

Genomically characterized pediatric neuroblastoma (NB) patient samples from the Therapeutically Applicable Research to Generate Effective Treatments (TARGET) study (Pugh et al., 2013; Wei et al., 2018) include 240 cases with high-risk NB, with patients between 1.5 and 16.5 years of age at diagnosis. The data was obtained from the Open Pediatric Cancer (OpenPedCan) project data release v12 (Geng et al., 2024), which is an open analysis effort to harmonize pediatric cancer data from multiple sources. We consider gene expression, mutation, and tumor purity, which is the proportion of cancer cells in a tumor sample compared to non-cancerous cells. Here we briefly summarize the data generating platforms and relevant data pre-processing used in this study. Gene expression was acquired using Illumina HiSeq 2500 platform and the measurements were reported in TPM values. The TPM gene expression data was *log*2 transformed with pseudo-count of 1. For mutation data, consensus calls from Strelka2 (Kim et al., 2018), Mutect2 (Benjamin et al., 2019), Lancet (Narzisi et al., 2018) and VarDictJava (Lai et al., 2016) were used. The mutation data was summarized with 1 indicating the presence of a mutation and 0 otherwise. Tumor purity was inferred using the THetA2 (Oesper et al., 2014) algorithm, which models somatic copy number aberrations, providing insights into the heterogeneity of the tumor. The values range from 0 (low-purity) to 1 (high-purity), where low values indicate higher fraction of non-cancerous cells (e.g., immune cells or stromal cells) relative to cancerous cells, while high values indicate predominance of cancerous cells in the tissue sample. We further excluded low variance genes (*σ* < 0.001) and missing data.

### 2.6 Neurobpastoma patient cohort from the Gabriella Miller Kids First (GMKF) Pediatric Research Program

The GMKF neuroblastoma Pediatric Research Program (Gabriella Miller Kids First Pediatric Research Program, 2017; Egolf et al., 2019) includes biospecimens from 1, 681 patients, who were between 0 to 5 years of age at diagnosis. We consider gene expression, mutation and tumor purity data from this cohort. The GMKF data was similarly obtained from the OpenPedCan project data release v12. This dataset followed identical processing steps as the TARGET cohort, described in section 2.5.

## 3 Methods

### 3.1 Notation and preliminaries

We consider *C* different cohorts, each with *M*_*c*_ different sets of measurements, which we will interchangeably call views: 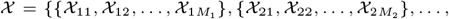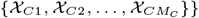. For each view 𝒳_*cm*_ with 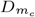 features and *N*_*c*_ observations, we assume an independently and identically distributed sample 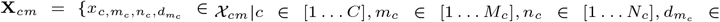 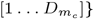, where *c, m*_*c*_, *n*_*c*_ and 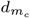 enumerate the cohorts, views, observations, and features, respectively. Generally, views can correspond to measurements by different biological assays (modalities), such as gene expression, copy number variation, mutation; functional perturbation assays, such as drug response; or measurements of the same modality acquired in different experiments. Views can have partially overlapping sets of features (Fig. 1). For example, omic modalities could profile shared subsets of genes, while functional assays could measure the cell viability of samples post drug treatment for the same subsets of drugs. We define a feature-wise prior 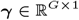 over the set of *G* unique features across all cohorts and views. Our model is generative and uses ***γ*** and view-specific projection matrices 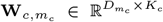 to learn a shared latent space representation simultaneously for each respective cohort 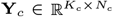 with *K*_*c*_ latent components (Fig. 2). We use 𝒩 (***µ*, Σ**) to denote the normal distribution with mean ***µ*** and covariance matrix **Σ**, and −(*α, β*) to denote the Gamma distribution with shape *α* and scale *β* hyperparameters. We use ⟨·⟩ to denote the expectation.

**Fig. 1.**
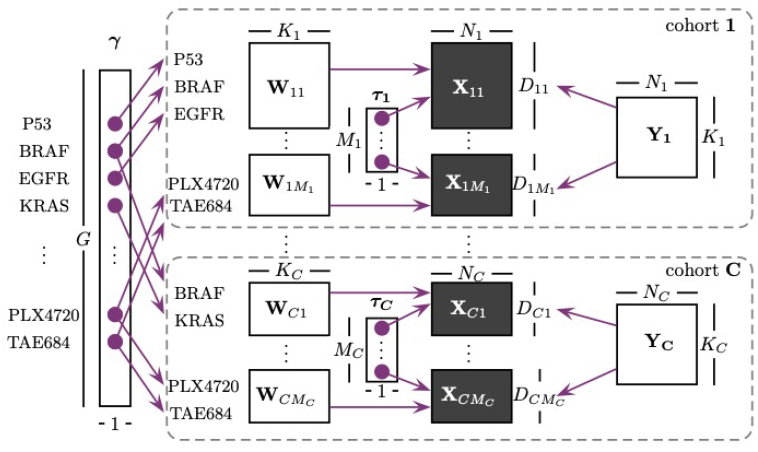
Matrix representation of the algorithm workflow illustrating dimensionality of variables and priors across *C* cohorts and their respective views. The feature-wise prior ***γ*** spans partially overlapping feature sets, including molecular features and drugs.

**Fig. 2.**
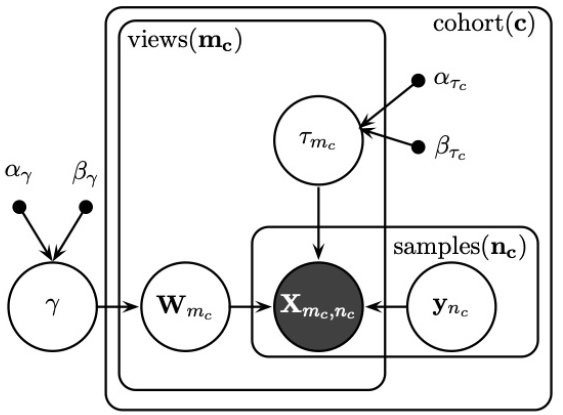
Graphical representation of the TLGFA model. The shaded node 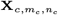 represents the observed variable and, the unshaded nodes of 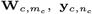 and 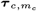 represent the latent variables. Arrows indicate causal relationships between variables and the rectangles represent the repeated structures. The parameter ***γ***, which is present outside the plate governs the distribution of 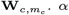 and *β* denote the shape and scale hyper-parameters of the gamma distribution, respectively.

### 3.2 Bayesian group factor analysis transfer learning (TLGFA) model

Graphical representation of the proposed model is shown in Fig. The joint likelihood of the TLGFA model is given by:

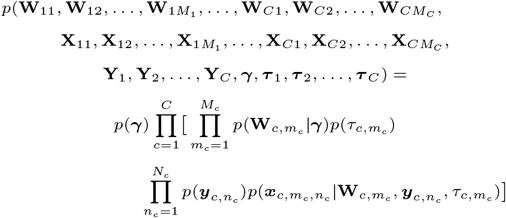

where,

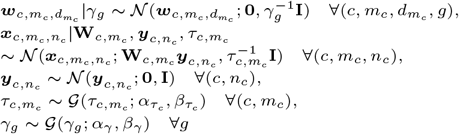

### 3.3 Inference

We present a deterministic variational approximation inference algorithm with closed-form update equations using conjugate priors. We approximate the full posterior via the following distribution and its factorization:

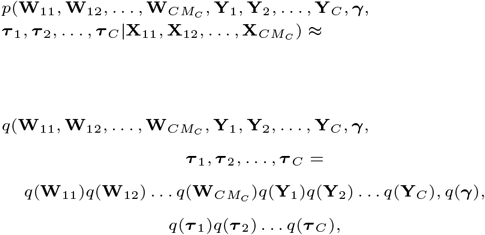

where each factor is defined to be from the family of its respective full conditional distribution. Recall that *G* denotes the total number of unique features across all views and cohorts. Then, the approximating distribution factorizes as follows:

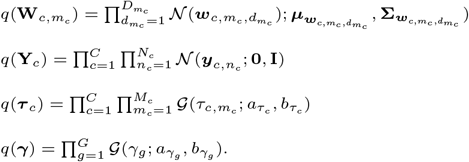

Minimizing the Kullback–Leibler divergence between the posterior 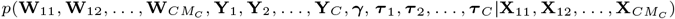 and the approximate posterior 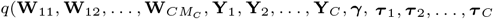 distributions is equivalent to maximizing the following lower bound derived using Jensen’s inequality:

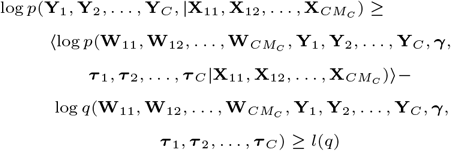

The algorithm optimizes the lower bound l(q) by updating each term of the bound until convergence. Let 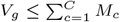 be the number of views in which the feature with index g in ***γ*** appears. Let *ξ*(*g, i, j*) be a many-to-one mapping function which returns the row 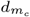 in projection matrix **W**_*ij*_ of the feature with index g in ***γ***.

Then the approximate posterior distribution for the feature-wise prior is given as follows:

Let 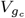 be the number of cohorts, and 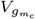 be the number of views in cohort c, such that the feature with index g in ***γ*** appears. That is, 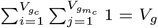 Then:

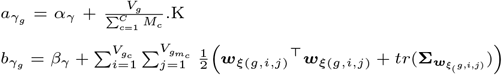

where, 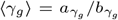.

The noise precision parameters update is given by:

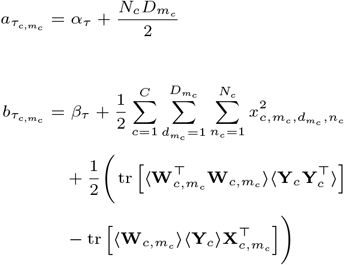

The latent variable’s approximate posterior is updated as follows:

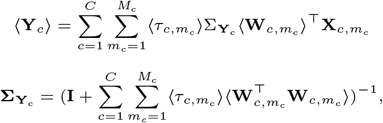

where,

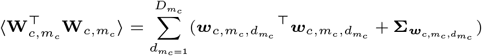

For each triplicate 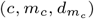 let 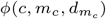 denote a many-to-one mapping function which returns the index g in the prior vector ***γ*** of the feature in row 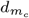 in projection matrix 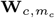 in cohort c. Then, the projections are updated as follows:

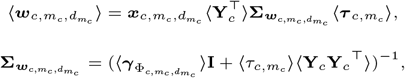

and

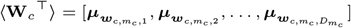

The lower bound is given in Supplementary File 1.

### 3.4 Prediction

Suppose that we want to predict views in a cohort 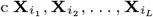, where *l* ∈ {1, …, *L*}, *i*_*l*_ ∈ {1, …, *M*_*c*_}, *L* < *M*_*c*_, given the views per cohort 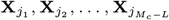, where {**X**_*i*_} and {**X**_*j*_} are mutually exclusive and 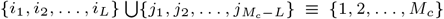. That is, we are now interested in the predictive distribution 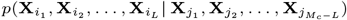. This distribution cannot be computed in closed form but can be derived using expectations from the variational approximation:

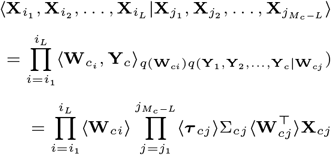

where,

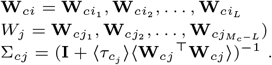

### 3.5 Comparison algorithms

We compare the performance of our proposed TLGFA model to several baseline models - the GBGFA (Nikolova et al., 2017), Elastic Net (ENET) (Zou and Hastie, 2005), Support Vector Regressor (SVR) (Vapnik, 1999; Pedregosa et al., 2011) using the Radial Basis Function kernel (Schölkopf et al., 1998), Random Forest Regressor (RFR) (Breiman, 2001; Pedregosa et al., 2011) and the Multi-Omics Factor Analysis (MOFA+) (Argelaguet et al., 2020).

GBGFA is a multitask Bayesian group factor analysis model similar to TLGFA, with a feature-wise prior that favors features that carry information in multiple measurement platforms. ENET is a linear regression method that uses lasso (*L*_1_) (Tibshirani, 1996) and ridge (*L*_2_) (Hoerl and Kennard, 1970) penalties to perform simultaneous feature selection and feature shrinkage. SVR is a robust machine learning method used to predict continuous values. It maps the input features to high-dimensional spaces to identify the ideal hyperplane that accurately fits the data. RFR is an ensemble-based supervised learning algorithm that constructs multiple decision trees from different data subsets and aggregates their outputs, to improve predictive performance. Finally, MOFA+ is a data integration framework which uses heterogeneous data to learn low-dimensional representations; while not suited for prediction, here we apply it to evaluate our performance in identifying predictive gene signatures.

## 4. Results

We evaluate our model’s performance on synthetic and primary cancer patient cohorts in acute myeloid leukemia (AML) and neuroblastoma (NB). First, we evaluate the model’s ability to predict drug response in AML patient samples by learning jointly from the BAML and FIMM cohorts. Second, we perform feature selection to assess our model’s ability to recapitulate consensus biomarkers of drug response. Next, we assess our model’s ability to predict tumor purity in NB using datasets from the TARGET and GMKF patient cohorts. Finally, investigate gene signatures of tumor purity learned by our model.

### 4.1 Synthetic data experiments

Here we evaluate if our model effectively encodes our underlying assumptions using controlled data generated in section 2.1. For a given feature shared across cohorts or views, we assume that if signal is present in one cohort, but undetectable on its own in another cohort, the result from joint training should be detectable, effectively implementing the boolean logic NOR operator. To assess the correctness of our approach under varying cross-domain conditions, we conducted a series of tests under distinct conditions: (1) features uninformative in both cohorts (`off-off` scenario), (2) features informative in both cohorts (`on-on` scenario), (3) features informative in the first cohort but uninformative in the second (`on-off` scenario) and (4) features uninformative in the first but informative in the second (`off-on` scenario). For each case, we selected 20 shared features from the first view of each cohort and set their corresponding factor loading vectors 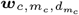 in 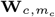. We set them to 1*e*–14 for uninformative (off) and to 1 for informative features (`on`) cases.

After training, we examined the factor loadings vectors for the 20 test features in each scenario to determine whether signal propagation occurred as expected. Results from the analysis are summarized in Fig. 3 for the `off-off` and on-off cases, and in supplementary figure SFig. 1 for the remaining cases. To quantify the signal distribution range, we computed the minimum and maximum values for the initial and learned factor loading vectors for the 20 tested features, using base R packages. Results from the analysis are summarized in Table 1.

**Table 1.** Comparison of minimum and maximum range of the initial and learned 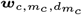 distributions for the 20 shared features for the four cases in both the cohorts.

| CASES | COHORT | MINIMUM (Initial) | MAXIMUM (Initial) | MINIMUM (Learned) | MAXIMUM (Learned) |
| --- | --- | --- | --- | --- | --- |
| Off - Off | 1 | $1e-14$ | $1e-14$ | $-2.65e-05$ | $2.27e-05$ |
| | 2 | $1e-14$ | $1e-14$ | $-2.06e-05$ | $2.49e-05$ |
| On - On | 1 | 1 | 1 | -0.938 | 1.190 |
|  | 2 | 1 | 1 | -1.207 | 1.202 |
| On - Off | 1 | 1 | 1 | -0.709 | 0.696 |
| | 2 | $1e-14$ | $1e-14$ | -0.027 | 0.028 |
| Off - On | 1 | $1e-14$ | $1e-14$ | -0.026 | 0.026 |
|  | 2 | 1 | 1 | -0.993 | 0.605 |

**Fig. 3.**
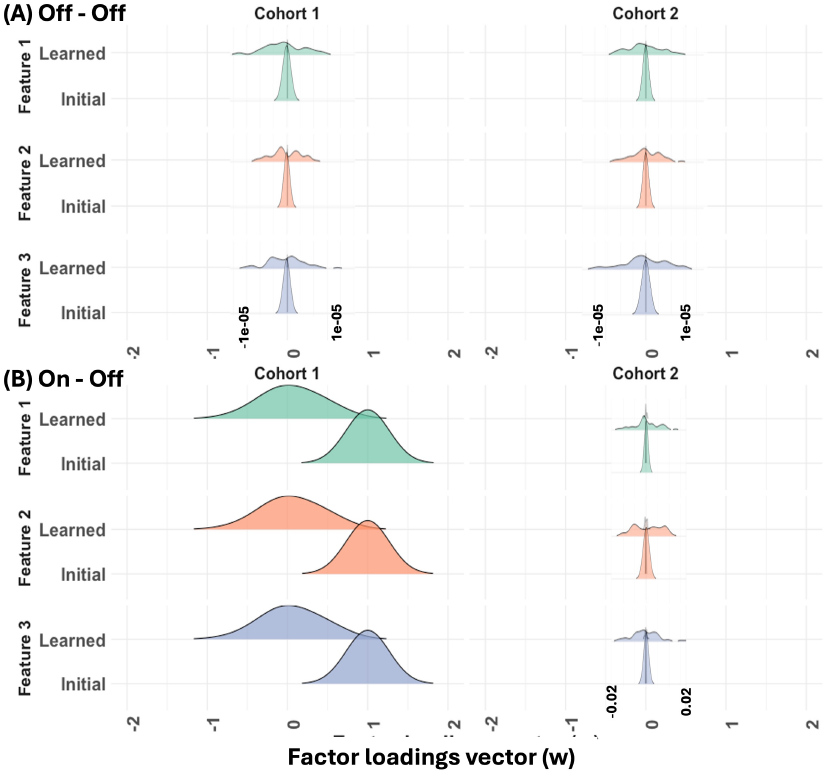
Ridge plots showing the distribution of factor loading vectors 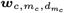 for three shared features across cohorts: (A) `Off-Off` and (B) On-Off. A distribution centered around 1*e* − 14 suggests low informativeness (off). This illustration allows us to visually assess how well the model preserves or transfers feature informativeness between cohorts. All plots are shown within the range of −2 to 2, with zoomed-in overlays highlighting finer details.

### 4.2 Performance evaluation of drug response prediction in AML

We evaluate our method’s performance in predicting drug response. We consider gene expression, mutation and drug response from the BAML and FIMM cohorts. We compare our model TLGFA to four established models, namely ENET, SVR, RFR, and GBGFA, as described in section 3.5 The TLGFA and GBGFA models were trained with *K*_*c*_ = 75 and *α*_*γ*_ = *β*_*γ*_ = *α*_*τ*_ = *β*_*τ*_ = 1*e* − 14. ENET, SVR and RFR were trained using sklearn python packages, with hyperparameter tuning performed through grid search method. ENET, SVR and RFR were each trained on one cohort and predictions were made for the other cohort. In contrast, TLGFA was trained jointly on both cohorts, and predictions were made using the corresponding projection matrices **W**_*c*_ for each cohort. GBGFA was trained separately on each cohort, with predictions made within the same cohort. The predictive accuracy is evaluated using 5-fold cross validation with the BAML and FIMM datasets. Each dataset is divided into five non-overlapping folds, with each fold used as a test set while the model is trained on the remaining four. The data was randomly divided into five partitions ten times to ensure that the results are independent of the partitioning process, and we report the average of the ten random splits. The baseline models are compared to TLGFA using the proportion of variance (POV) metric, defined as the complement of the normalized root mean square error (NRMSE): POV = 1-NRMSE,

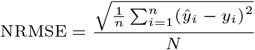

where *ŷ*_*i*_ and *y*_*i*_ are predicted and observed values, respectively. The TLGFA and GBGFA models were trained on shared and distinct drugs. However, due to the limitation of ENET, SVR and RFR to predict only on drugs shared across both the cohorts, we considered only the shared drugs (53 drugs were shared across both the cohorts) for further analysis. The results of these experiments are summarized in Fig. 4. Paired Wilcoxon test *p*-value is also reported for each comparison.

**Fig. 4.**
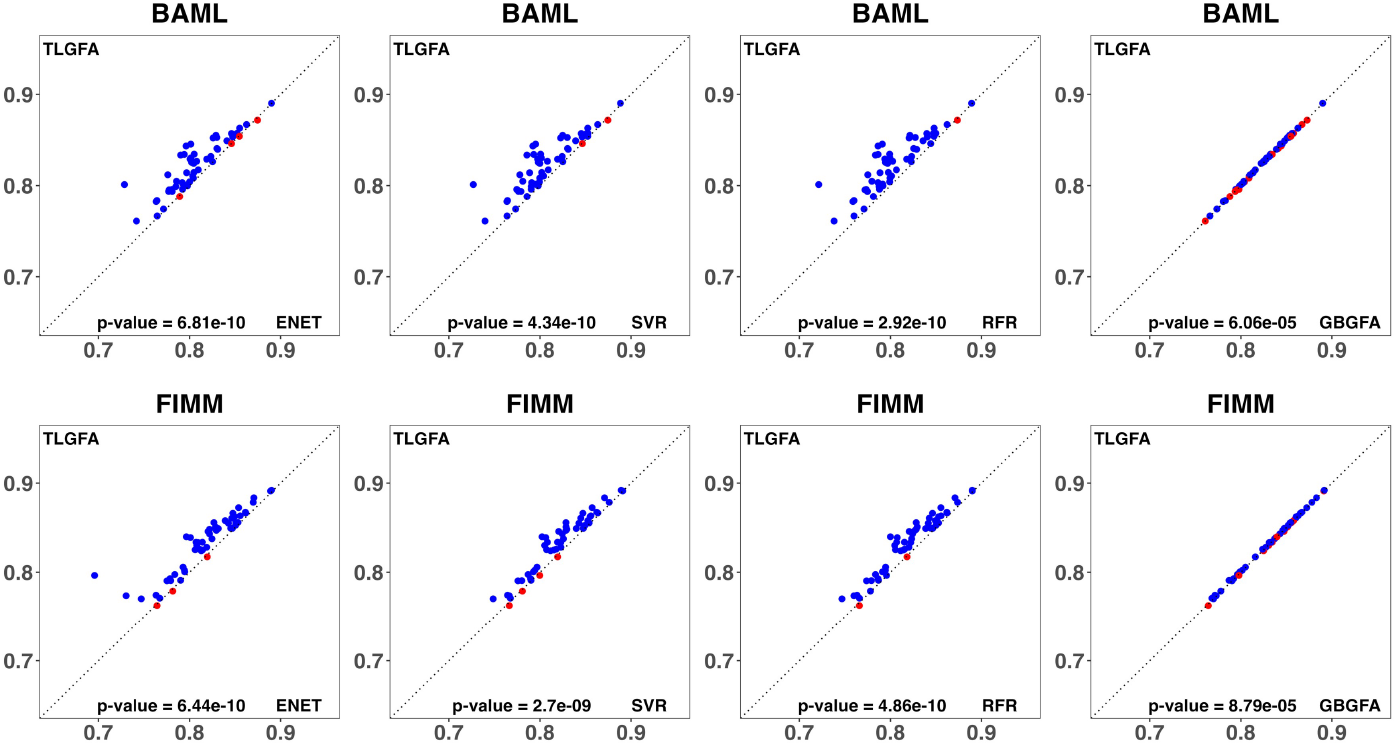
Cross-validation results of drug response prediction in AML. The top and bottom rows correspond to the BAML and the FIMM panels, respectively. Dot-plots present in columns 2 through 5 shows the pairwise comparison of TLGFA (Y-axis) and Elastic Net (X-axis), TLGFA (Y-axis) and Support vector regressor (X-axis), TLGFA (Y-axis) and Random forest regressor (X-axis), and TLGFA (Y-axis) and GBGFA (X-axis), respectively, where each dot shows the POV for individual shared drugs between the two cohorts. Significance was evaluated through paired Wilcoxon test.

### 4.3 Performance evaluation of consensus biomarkers recovery in AML

We examine the consensus biomarkers of drug response which were curated and summarized in supp. table ST1 assembled as described in section 2.4. We compare the results of TLGFA with two baseline models - GBGFA and MOFA+. All models were trained on gene expression, mutation and drug response data from BAML and FIMM. For each model, we tested three different *K* values and present the best out of 3 results, yielding *K*_*c*_ = 75 for TLGFA, GBGFA, and *K*_*c*_ = 20 for MOFA+. For both cohorts, only drugs shared across cohorts, with a win in the respective cohort, for which consensus biomarkers were available were considered. We calculate gene scores for each consensus biomarkers as follows. For each drug *i* and view *m*, we first calculate gene scores 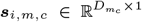, as the product of the view-weights and drug response vector 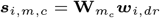 in every cohort *c*, where *dr* denotes a drug response matrix in either BAML or FIMM. We calculate the cumulative density function (CDF) using the kernel density as input to the R package Spatstat. We compute a final gene score per cohort by optimizing values on both tails of view-specific gene CDFs: 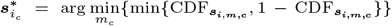. Lower CDF values indicate strong recapitulation ability. We summarize the performance per drug by taking the median gene score of all available consensus biomarkers for the given drug. We summarize our performance by taking the difference ∆ = baseline_*i*_− TLGFA_*i*_ of median gene scores for each drug *i*. Positive ∆ values indicate improvement over baseline approaches (win), negative - otherwise (loss). The results of these experiments are shown in Fig. 5.

**Fig. 5.**
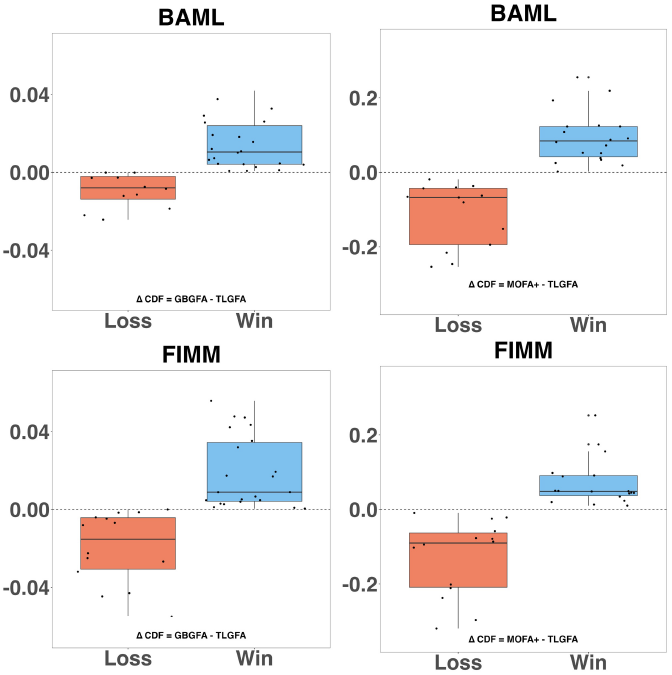
Consensus biomarker analysis results. The top and bottom rows correspond to BAML and FIMM, respectively. Left column compares TLGFA to GBGFA, and column right column compares TLGFA and MOFA+, respectively. Each dot in the boxplots represents the difference ∆ between the median CDF values of the biomarkers for each drug (Y-axis). Blue and red boxplots correspond to drugs in the win and loss cases.

We further annotated drugs with their corresponding families based on classifications from Bottomly et al, supp. file 4 (Bottomly et al., 2022) (section 2.4), and the wins and losses were analyzed at the drug family level. The results of this analysis comparing the TLGFA to GBGFA and MOFA+ are summarized in supplementary figures SFig. 2 and SFig. 3, respectively.

### 4.4 Performance evaluation of tumor purity prediction in NB

We evaluate our model’s ability to predict tumor purity in two neuroblastoma patient cohorts. Gene expression, mutation and tumor purity were considered from the two cohorts. The predictive performance of TLGFA was compared with four established models as before, ENET, SVR, RFR, and GBGFA. The TLGFA and GBGFA models were initialized with *α*_*γ*_ = *β*_*γ*_ = *α*_*τ*_ = *β*_*τ*_ = 1*e* − 14, and trained with *K*_*c*_ = 50. The rest of the models were trained using sklearn python packages and grid search was used for hyperparameter tuning. TLGFA was trained across both cohorts simultaneously, leveraging the respective projection matrices **W**_*c*_ for prediction. GBGFA, on the other hand, was trained independently on each cohort, with predictions restricted to the same cohort it was trained on. ENET, SVR, and RFR models were trained on a single cohort and then used to make predictions on the other. To evaluate the predictive accuracy of the models, we perform 5-fold cross-validation with 10 random splits into folds for each dataset. We assess performance using the POV and the mean absolute error (MAE) metrics, where

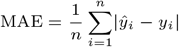

where *ŷ*_*i*_ and *y*_*i*_ are predicted and observed values, respectively. The results of these experiments are summarized in Table 2.

**Table 2.**
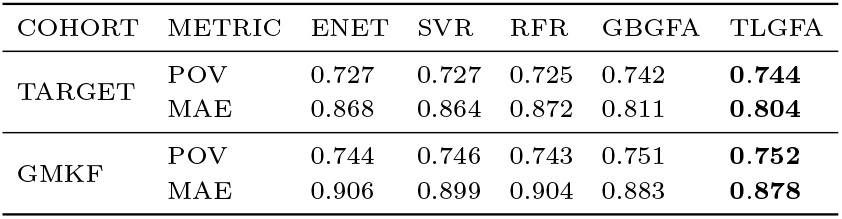
Cross-validation results for tumor purity prediction in neuroblastoma. Each cohort shows both the Proportion of Variance (POV) and Mean Absolute Error (MAE). The models with the highest POV and lowest MAE are marked in bold for each cohort.

| COHORT | METRIC | ENET | SVR | RFR | GBGFA | TLGFA |
| --- | --- | --- | --- | --- | --- | --- |
| TARGET | POV | 0.727 | 0.727 | 0.725 | 0.742 | **0.744** |
|  | MAE | 0.868 | 0.864 | 0.872 | 0.811 | **0.804** |
| GMKF | POV | 0.744 | 0.746 | 0.743 | 0.751 | **0.752** |
|  | MAE | 0.906 | 0.899 | 0.904 | 0.883 | **0.878** |

#### 4.4.1 Tumor purity signatures identification in NB

Neuroblastoma is a pediatric solid tumor that originates from the developing peripheral sympathetic nervous system with median age of diagnosis at 17 months (London et al., 2005). The tumor presents with heterogeneity in clinical phenotype, localization, and genetic abnormalities (Mahapatra and Challagundla, 2025). One crucial factor that leads to tumor heterogeneity is the influence of the tumor microenvironment (Gomez et al., 2022). A key aspect of this landscape is tumor purity which can potentially influence molecular profiling, therapeutic response, and patient outcomes. Gene signatures associated with tumor purity can help delineate some of these signals. Here we derive a gene signature predictive of tumor purity. First, we calculate gene scores 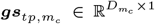 as the product of the view-specific weights and 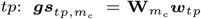 in every cohort *c* followed by Z-score standardization, where tumor purity *tp* is either from the TARGET or the GMKF cohorts. An overall gene score for tumor purity and gene *g* is computed by optimizing the absolute value of view-specific standardized gene scores 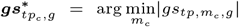. The computed gene scores are used to rank genes in decreasing order of importance, with the lowest rank being the most relevant. We select the top 50 ranked genes. The inferred signature is projected back onto the genomic training data, where genes are ordered by their respective gene scores and patient samples are ordered by tumor purity values. For ease of presentation here we highlight the 25% of patient samples with the highest and lowest tumor purity, respectfully (*N* = 21). Results for the TARGET cohort are summarized in Fig. 6 and for the GMKF cohort are summarized in supplementary figure SFig. 4.

**Fig. 6.**
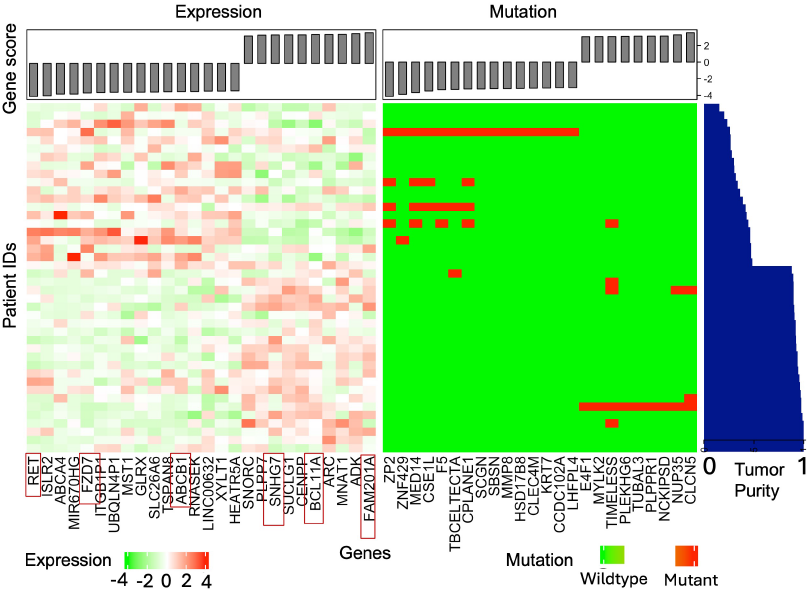
Predictive gene signature analysis in the TARGET patient cohort. The gene signature from gene expression and mutation is projected back onto the genomic training data, where genes (X-axis) are ordered based on their respective model score (top grey bar) and patient samples (Y-axis) are ordered based on tumor purity levels. Genes with known associations with neuroblastoma highlighted in red.

## 5. Discussion

### Proposed model encodes intended behavior

We first assessed if our model effectively encodes our intent, namely that for any given feature shared across cohorts or modalities, we assume that if signal is present in one cohort, but undetectable on its own in another cohort, the result from joint training should be detectable, effectively implementing the boolean logic NOR operator. We examined this by comparing the initial and learned values of the factor loading vectors 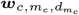 for 20 synthetic features in 4 key scenarios described in section 4.1. In the `off-off` case, where shared features were undetectable in both cohorts (Fig. 3 panel A), the range of initial values (minimum and maximum 1*e*−14) remained similar after training, where learned values ranged from −2.65*e* − 05 to 2.5*e* − 05 as shown in the first row of Table 1. This outcome aligns with our expectation, confirming that in the absence of detectable signal across all cohorts, the model does not artificially introduce signal. In contrast, for the on-on case, where shared features were detected in both domains (supplementary figure SFig. 1 panel A), the learned factor loadings remained similar to their initialization, where initial values: min = max = 1; learned values: cohort 1 min = −0.938, max = 1.190; cohort 2 min = −1.207, max = 1.202 as shown in the second row of Table 1. This indicates that TLGFA preserves detectable signals when present in both domains. In the on-off case, cohort 1 began with detectable values of min = max = 1, while cohort 2 started undetectable min = max = 1*e* − 14 (Fig. 3 panel B). After training, cohort 2’s factor loadings shifted to a distribution closer to the an on case with a min = −0.027 and a max = 0.028, indicating successful signal transfer, as shown in row 3 of Table 1. A similar trend was observed in the off-on case (supplementary figure SFig. 1 panel B), where cohort 1 recovered signal after training (min = −0.026, max = 0.026) despite starting with no signal (refer to Table 1, row 4). Collectively, these results confirm that TLGFA effectively transfers information across domains in all three information-positive scenarios, without introducing noise in the fully undetectable setting. This performance supports the intended effect of the feature-wise prior enabling the model to corroborate positive signal.

### Proposed model improves drug response prediction across independent AML patient cohorts

We analyze the predictive performance of TLGFA using data from two AML patient cohorts. We compare the performance of TLGFA with four existing models as described in section 4.2. Using 5-fold cross-validation and ten random data splits, we report the POV for each drug and summarize the results in Fig. 4. In the BAML cohort, TLGFA performed significantly better than ENET (*p*-value of 6.81*e* − 10), SVR (*p*-value of 4.34*e* − 10), RFR (*p*-value of 2.92*e* − 10) and GBGFA (*p*-value of 6.06*e* − 05). We observed a similar trend the FIMM cohort: ENET (*p*-value of 6.44*e* − 10), SVR (*p*-value of 2.7*e* − 09), RFR (*p*-value of 4.86*e* − 10) and GBGFA (*p*-value of 8.79*e* − 05). Our comparison reveals statistically significant improvements in drug prediction across both cohorts. Consistent with previous results, multi-task models like GBGFA and TLGFA outperform single-task models, highlighting the advantages of multi-tasking. In summary, our results indicate that our feature-wise prior facilitates cross-cohort information transfer and improves predictive accuracy compared to the established models in both the cohorts.

### Proposed model improves identification of consensus biomarkers

Drug signatures, where a set of genes is identified as either predictive of positive drug response or drug resistance, are of interest for multiple reasons: (i) genes identified to be associated with resistance could help prioritize novel drug targets; and (ii) transcriptional signatures in particular could be used to stratify treatment for patients who do not present with a known driver mutation, and thus do not fall into an established genotype, which is a common occurance in AML. In our assessment of our proposed model to recapitulate drug signatures, we focus in particular on established consensus biomarkers of drug response, when those are available (Supp. table ST1). Our results from comparing TLGFA with two existing models as described in section 4.3 are summarized in Fig. 5. Our results indicate an overall trend of improved biomarker ranking in TLGFA as compared to the existing approaches. When compared to GBGFA, TLGFA outperformed in 22 of 34 drugs in BAML, and 22 of 37 drugs in FIMM (Fig. 5, left column). A similar trend, although not as pronounced, was also observed when comparing to MOFA+. Comparing the results from TLGFA and GBGFA, in the BAML cohort, most of the drugs with improved performance belonged to the Type III and IV Receptor Tyrosine Kinase (RTK) family showed positive outcomes (supplementary figure SFig. 2 panel A), while in the FIMM cohort, RTK inhibitors and STE family generally exhibited a higher number of wins compared to other drug families (refer to supplementary figure SFig. 2 panel B). We observed a similar trend with TLGFA vs MOFA+ (supplementary figure SFig. 3). In summary, we observed that the transfer learning approach is able to improve biomarker ranking while learning from two geographically diverse patient populations of the USA and Finland.

### Proposed model improves prediction accuracy of tumor purity prediction in NB patient cohorts

Tumor purity is a critical variable to consider, as solid tumor tissues contain both cancerous and non-cancerous cells and these non-cancerous cells have been associated with tumor progression and poor clinical outcomes in several cancers (Sato et al., 2005; Mlecnik et al., 2011). Accurately estimating tumor purity may provide insights into the tumor microenvironment and its role in patient prognosis. TARGET and GMKF consist of diverse patient samples making them well-suited case study for our method.

We assess TLGFA’s ability to predict tumor purity by simultaneously learning from two NB patient cohorts with distinct population profiles. Low purity samples, characterized by a higher proportion of non-cancerous cells, can obscure tumor-specific molecular signals, making biomarker detection and molecular sub-typing more difficult. As a result, accurate tumor purity estimation can aid genomic analysis, biomarker discovery, and treatment planning. Predictive performance of TLGFA was compared with four baseline models as described in section 4.4. We report the POV and MAE for each cohort using 5-fold cross-validation and 10 random data splits and summarized the results in Table 2. We next highlight the results in pairs, where in each result pair, the first value indicates POV and the second indicates MAE. In the TARGET cohort, TLGFA (0.744, 0.804) outperforms ENET (0.727, 0.868), SVR (0.727, 0.864), RFR (0.725, 0.872), and GBGFA (0.742, 0.811). A similar trend is observed in the GMKF cohort, where TLGFA (0.752, 0.878) again surpasses ENET (0.744, 0.906), SVR (0.746, 0.899), RFR (0.743, 0.904), and GBGFA (0.751, 0.883). Overall, our results indicate that our model improves prediction accuracy in both cohorts.

### Proposed model identifies gene signatures associated with different tumor purity levels

Gene signatures associated with varying tumor purity levels can be identified using the interpretable latent space representations learned by TLGFA. We also assess the benefit of incorporating a feature-wise prior in uncovering these gene signatures. In the TARGET cohort, we identified multiple genes directly linked to neuroblastoma, including ADK, BCL11A, and SNHG7, which are typically up-regulated in this cancer (Chen et al., 2019; Jia et al., 2020; Martínez-Pacheco et al., 2023). Loss of expression in these genes has been related to inhibition of cell growth and migration, as well as the initiation of apoptosis (Fig. 6 - highlighted in red). These genes appear to be tied to high tumor purity in our analysis. In contrast, FZD7, another gene commonly up-regulated in neuroblastoma, and whose inhibition may trigger apoptosis (Ahmad et al., 2023), is found to be correlated with low tumor purity (Fig. 6 - highlighted in red). TLGFA also recovered FAM201A (Ye et al., 2023) and RET (Siaw et al., 2021), whose loss is known to contribute to proliferation and metastasis in neuroblastoma. In our analysis, up-regulated RET and down-regulated FAM201A are attributed to lower tumor purity, suggesting a complex regulatory role. Another key gene identified is ABCB1, previously linked to drug resistance (Fig. 6 - highlighted in red) in neuroblastoma (Löschmann et al., 2016). In the GMKF cohort, TLGFA recovered L3MBTL2, a gene shown to interact with MYCN to sustain cell proliferation

(Okada et al., 2024). Its up-regulation is connected with low tumor purity in our analysis (refer to supplementary figure SFig. 4 - highlighted in red). TLGFA also uncovered additional genes such as CSE1L (Li et al., 2024), F5 (Liu et al., 2020), TACC2 (Onodera et al., 2016) and ZFP36L2 (Lin et al., 2025), which have been linked with poor prognosis in other cancers. While these findings are empirical and need further biological validation, our results indicate that the feature-centric prior improves prioritization of driver genes that corroborate existing literature in both cohorts. It helps refine known biomarkers of neuroblastoma and identifies additional downstream markers which further deconvolute characteristics of samples with lower tumor purity levels vs. high tumor purity.

## Conclusion

We present a novel transfer learning approach, with fully Bayesian formulation and multi-task capabilities, which enables for the simultaneous knowledge transfer across disjoint, heterogeneous domains while incorporating feature-wise dependencies. We evaluate our approach on both synthetic data and primary patient cancer datasets from acute myeloid leukemia (AML) and pediatric neuroblastoma (NB). We demonstrate the model’s effectiveness in predicting drug response in AML and tumor purity in NB, two clinically relevant regression tasks. Our results indicate that our feature-wise prior facilitates the transfer of biologically relevant information and enhances the prediction accuracy across multiple cohorts. The model also improves feature selection evaluated via the application to recapitulate known biomarkers of drug response. Overall, our proposed framework offers a scalable, generalizable and interpretable solution for leveraging existing datasets through joint multi-cohort learning.

## Data availability

The datasets used in this study are publicly available and described in section 2 and cited throughout this paper. Drug annotations with biomarkers is provided in the supp. file SF1.

## Code availability

The code for this work is available at https://gitlab.com/oats2/tl_gbgfa.

## Acknowledgments

We would like to acknowledge Liye He, Ph.D. and Caroline Heckman, Ph.D. from the University of Helsinki, Finland, for providing the drug response data for the FIMM cohort.

This work was partially funded by NIH NCI K22CA258799 computational infrastructure used was supported by NIH S10OD034224.

