## Supplementary figures for "Transfer learning framework via Bayesian group factor analysis incorporating feature-wise dependencies"

D Thirumalaisamy, et al.

### 1 Lower bounds

The lower bound is given by:

$$\begin{aligned}\mathcal{L}(q) = &\sum_{c=1}^{C}\sum_{m_c=1}^{M_c}\sum_{n_c=1}^{N_c}\log\langle p(\boldsymbol{x}_{c,m_c,n_c}|\mathbf{W}_{c,m_c},\boldsymbol{y}_{c,n_c},\tau_{c,m_c})\rangle \\ &+\sum_{g=1}^{G}(\log\langle p(\gamma_g)\rangle - \log\langle q(\gamma_g)\rangle) \\ &+\sum_{c=1}^{C}\sum_{m_c=1}^{M_c}(\log\langle p(\tau_{c,m_c})\rangle - \log\langle q(\tau_{c,m_c})\rangle) \\ &+\sum_{c=1}^{C}\sum_{m_c=1}^{M_c}\sum_{d_{m_c}=1}^{D_{m_c}}\big(\log\langle p(\boldsymbol{w}_{c,m_c,d_{m_c}}|\gamma_{\phi(c,m_c,d_{m_c})})\rangle \\ &-\log\langle q(\boldsymbol{w}_{c,m_c,d_{m_c}})\rangle\big) \\ &+\sum_{c=1}^{C}\sum_{n_c=1}^{N_c}\big(\log\langle p(\boldsymbol{y}_{c,n_c})\rangle - \log\langle q(\boldsymbol{y}_{c,n_c})\rangle\big)\end{aligned}$$

where $\langle . \rangle$ denotes expectation under $q$.

### 2 Supplementary figures

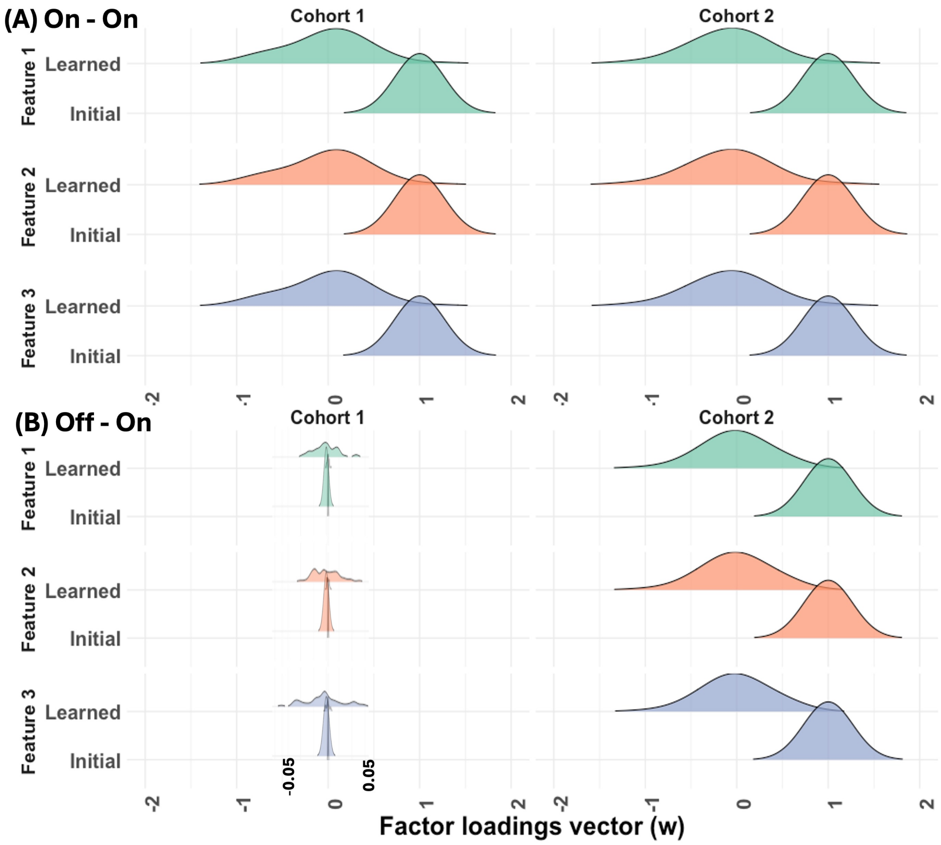


**Fig. S1.** Ridge plots showing the distribution of factor loading vectors $\boldsymbol{w}_{c,m_c,d_{m_c}}$ for three shared features across cohorts: (A)`On-On`, and (B) `Off-On` cases. A distribution centered around $1e-14$ suggests low informativeness (off). This allows us to visually assess how well the model preserves or transfers feature informativeness between cohorts. All plots are shown within the range of $-2$ to 2, with zoomed-in overlays highlighting finer details.

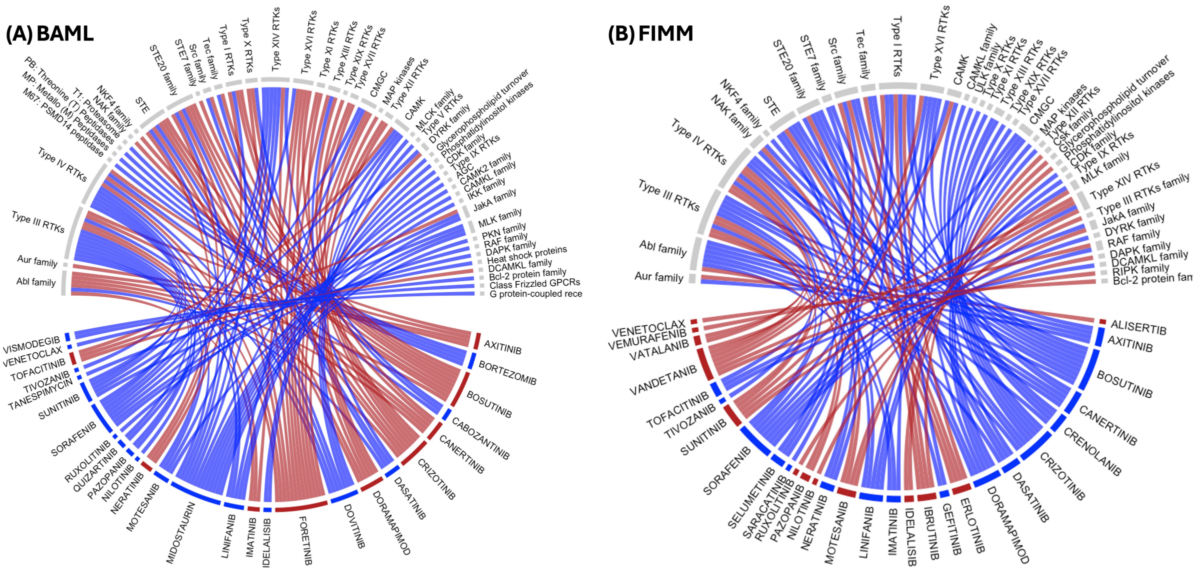


**Fig. S2.** Circos plots illustrating drug and drug family annotations outcomes for the TLGFA vs. GBGFA comparison. Wins (blue curves) and losses (red curves) are shown per drug family for (A) the BAML dataset and (B) the FIMM dataset. Each curve represents a single drug, and the total number of curves linked to a drug family indicates the number of drugs belonging to that family.

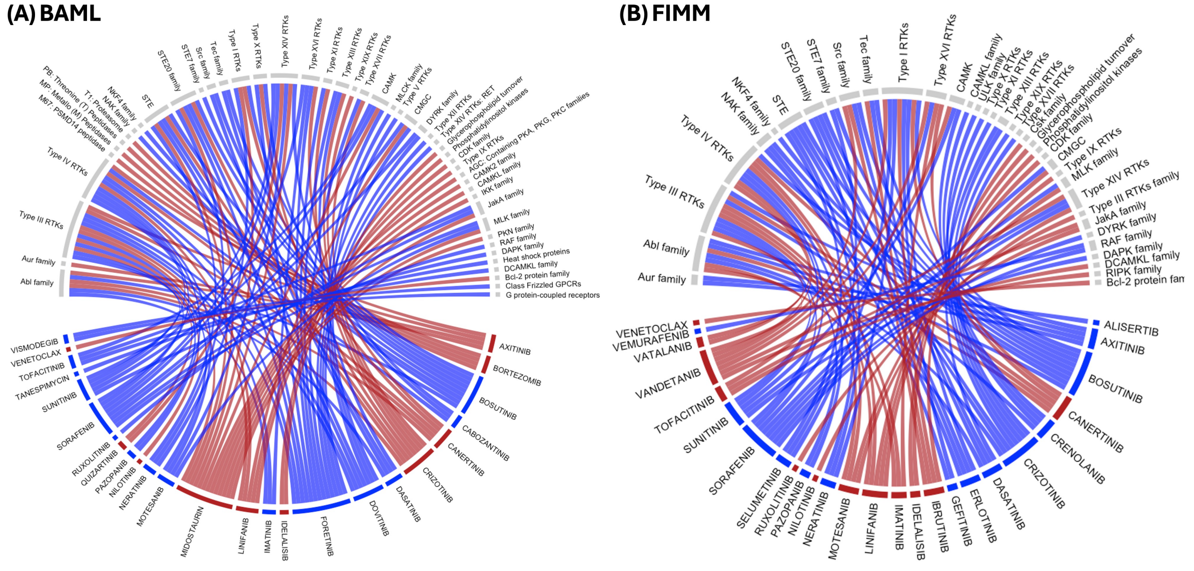


**Fig. S3.** Circos plots comparing TLGFA and MOFA+ show drugs and drug family annotation results. Blue curves indicate wins and red curves indicate losses for each drug family in (A) the BAML and (B) the FIMM datasets. Each curve corresponds to one drug, and the number of curves per family reflects the total drugs in that category.

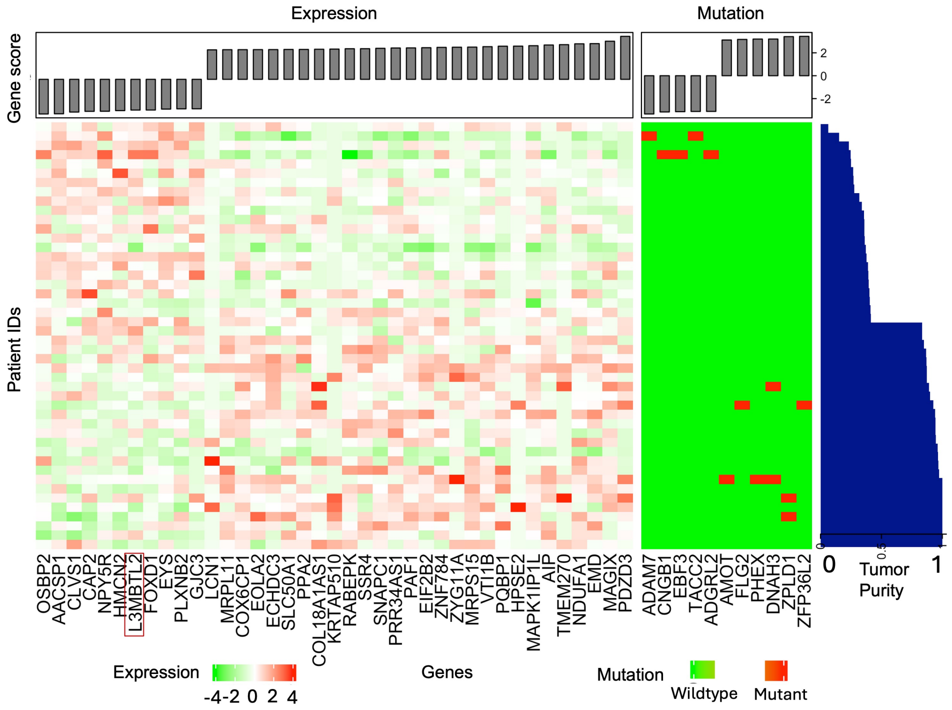


**Fig. S4.** Predictive gene signature analysis in the GMKF cohort. The inferred signature from model trained on GMKF cohort's expression and mutation is projected back onto the genomic training data, where genes (X-axis) are ordered based on their respective model score (top grey bar) and patient samples (Y-axis) are ordered based on tumor purity levels. The model clearly distinguished gene signatures predictive of low and high purity. Genes mutated in the low purity are wild type in the high purity and vice versa. Similarly, genes up-regulated in the low purity are down-regulated in high purity and vice versa. Genes with known associations with neuroblastoma highlighted in red.
